# First-in-human demonstration of floating EMG sensors and stimulators wirelessly powered and operated by volume conduction

**DOI:** 10.1101/2023.06.30.547056

**Authors:** Laura Becerra-Fajardo, Jesus Minguillon, Marc Oliver Krob, Camila Rodrigues, Miguel González- Sánchez, Álvaro Megía-García, Carolina Redondo Galán, Francisco Gutiérrez Henares, Albert Comerma, Antonio J. del-Ama, Angel Gil-Agudo, Francisco Grandas, Andreas Schneider-Ickert, Filipe Oliveira Barroso, Antoni Ivorra

## Abstract

**Background:** Recently we reported the design and evaluation of floating semi-implantable devices that receive power from and bidirectionally communicate with an external system using coupling by volume conduction. The approach, of which the semi-implantable devices are proof-of-concept prototypes, may overcome some limitations presented by existing neuroprostheses, especially those related to implant size and deployment, as the implants avoid bulky components and can be developed as threadlike devices. Here, it is reported the first-in-human acute demonstration of these devices for electromyography (EMG) sensing and electrical stimulation.

**Methods:** A proof-of-concept device, consisting of implantable thin-film electrodes and a non-implantable miniature electronic circuit connected to them, was deployed in the upper or lower limb of six healthy participants. Two external electrodes were strapped around the limb and were connected to the external system which delivered high frequency current bursts. Within these bursts, 13 commands were modulated to communicate with the implant.

**Results:** Four devices were deployed in the biceps brachii and the gastrocnemius medialis muscles, and the external system was able to power and communicate with them. Limitations regarding insertion and communication speed are reported. Sensing and stimulation parameters were configured from the external system. In one participant, electrical stimulation and EMG acquisition assays were performed, demonstrating the feasibility of the approach to power and communicate with the floating device.

**Conclusions:** This is the first-in-human demonstration of EMG sensors and electrical stimulators powered and operated by volume conduction. These proof-of-concept devices can be miniaturized using current microelectronic technologies, enabling fully implantable networked neuroprosthetics.

## I. Background

Neuroprosthetic technologies based on active implantable medical devices (AIMDs) have the potential to be an important tool for improving the quality of life of patients with neurological disorders and patients that require external systems as prostheses and exoskeletons. For example, intramuscular microstimulators could be used to reduce pathological tremor in essential tremor patients, as intramuscular electrical stimulation has been shown to reduce tremor in this population of patients up to 24 h (1). Moreover, intramuscular electromyography (EMG) microsensors could be used to control prostheses and exoskeletons (2–4), especially as it has been demonstrated to be a reliable input to feed neuromusculoskeletal models to estimate the user-intended joint movements for human-machine interface (HMI) control (5).

Implantable neuroprostheses form a chronic interface with the nervous system that overcomes one of the most important limitations of EMG sensors and electrical stimulators based on surface electrodes: the lack of selectivity. In addition, implantable electrodes improve repeatability by avoiding the impact of skin impedance changes and movements, and allow accessing deep muscles. In the case of EMG sensing for HMI, implantable electrodes avoid the need to replace the electrodes at each use (which in turn requires recalibration of the HMI) (6) and crosstalk, and allow the control of more degrees-of-freedom through the use of multiple electrodes (7).

Within the EXTEND collaborative project, funded by the European Commission and in which the authors of this study participated, it has been defined the concept of Bidirectional Hyper-Connected Neural Systems (BHNS). It refers to systems consisting of minimally invasive communication links between multiple nerves or muscles in the body and external devices which may be digitally interconnected between them (Fig. 1a). This will provide the means of a synthetic chain of action-reaction of sensorimotor activity using musculoskeletal modelling, aiding in applications as tremor management and HMI control. The BHNS is envisioned as a dense network of wireless implantable devices that can act as stimulators and EMG sensors, and that communicate in real time with the external systems. The external systems, in turn, process and analyze the neuromuscular activity and control the stimulation (e.g., for tremor management) and/or the action of machines (e.g., exoskeletons).

**Fig. 1.**
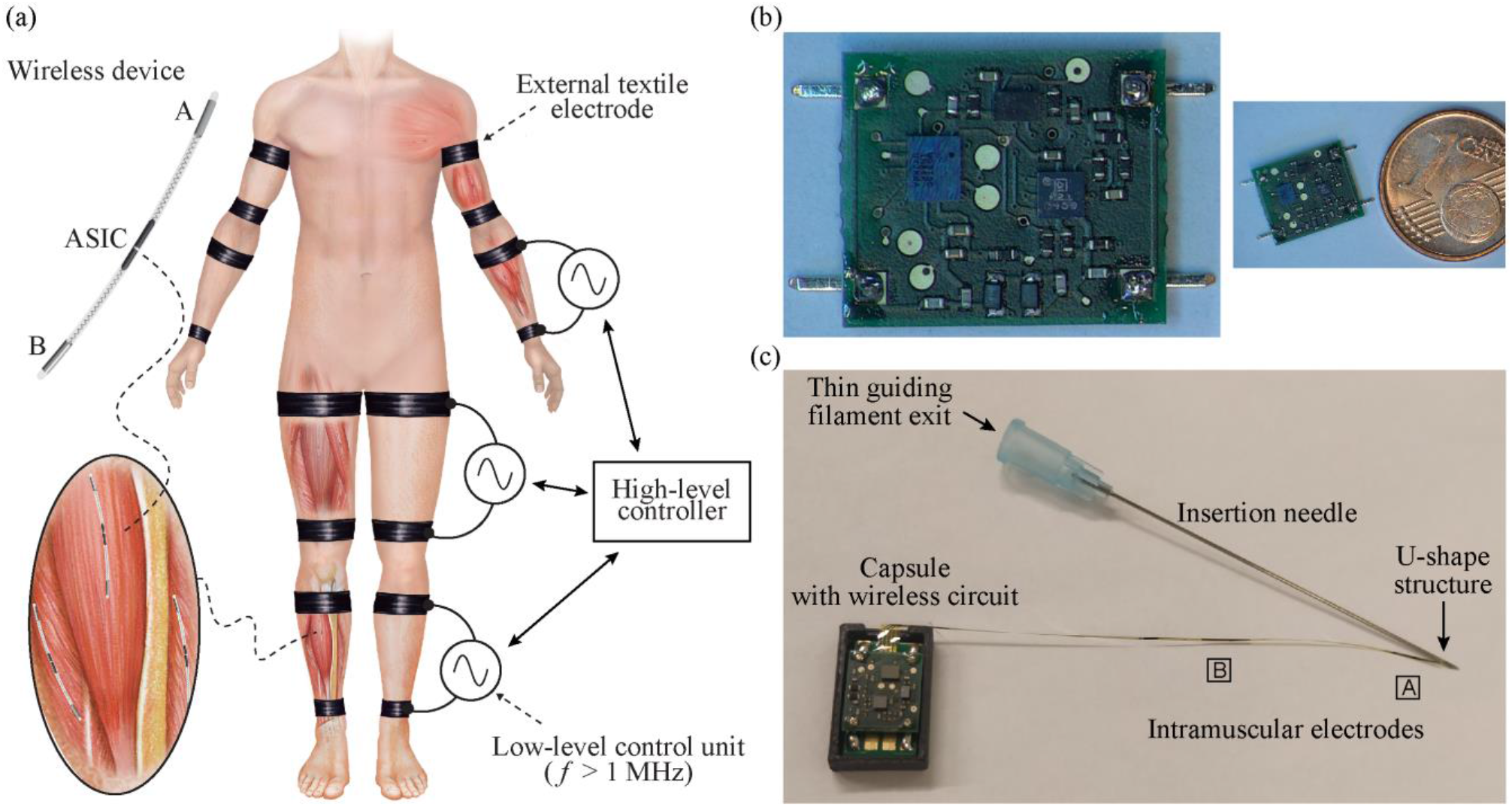
Bidirectional Hyper-Connected Neural System (BHNS) concept and proof-of-concept semi-implantable devices. a) Schematic representation of the envisioned BHNS. b) Miniature electronic circuit for EMG sensing and stimulation and its comparison with a 1 cent euro coin for acute human demonstration of the BHNS concept. c) Complete proof-of-concept semi-implantable device showing the dedicated insertion needle, the thin-film intramuscular electrodes, the thin guiding filament that exits through the needle hub, and the polymer capsule that houses the miniature circuit (opened lid) (17).

This BHNS paradigm requires fully implantable wireless devices that can be easily deployed by injection. Percutaneous systems would not be adequate to implement the BHNS concept because they require daily maintenance and dressing, and may present sweat gland blockage (7). Most fully implantable devices are developed as a central metallic case that houses the control unit, power circuit and pulse generator. The electronics are connected to the electrodes through leads, requiring complex surgical implantation procedures. Moreover, the leads tend to fail (8,9). As with the percutaneous systems, this approach would not be adequate for the implementation of the BHNS concept. To avoid the leads and their associated complications, it has been proposed the development of wireless networks of single-channel injectable devices that integrate the electrodes and the electronics. Nevertheless, the injectable devices capable of intramuscular stimulation that were developed in the past are stiff and relatively large (diameter > 2 mm) due to the need of batteries — with their limited lifespan and large volume (10) — or to the need of components required for the wireless power transfer (WPT) method used to energize them (e.g., coils in inductive coupling (11–13)). Recently, WPT methods as inductive coupling, ultrasonic acoustic coupling and capacitive coupling have obtained high miniaturization levels at the expense of link efficiency, penetration depth, or functionality. Reviews on these methods can be found in (10,14–16).

In the last years we have proposed and demonstrated a WPT method that avoids the use of bulky components inside the implant, and that allows to implement the implants as thin, flexible and elongated devices suitable for implantation by means of injection (18,19). The implants are powered and controlled by applying — through textile electrodes — high frequency (HF) current bursts that flow through the tissues by volume conduction (20). The currents are picked-up by the implant’s electrodes (located at the ends of the elongated body, Fig. 1a) and are rectified for powering and bidirectional communications. This allows the integration of the electronics in an application-specific integrated circuit (ASIC). We have validated in humans that HF current bursts complying with safety standards and applied through two textile electrodes strapped around a limb can provide substantial power from pairs of implanted electrodes, and are innocuous and imperceptible (21).

We have recently reported the development and evaluation of semi-implantable devices based on WPT by volume conduction that are capable both of performing stimulation and EMG sensing (17). These are proof-of-concept prototypes of the electronic architecture that will be integrated into an ASIC to obtain flexible threadlike devices as those demonstrated *in vivo* in (22), with stimulation, EMG sensing and bidirectional communication capabilities. The external system and a bidirectional communications protocol that allows wireless communication between the external system and the semi-implantable devices are also described in (17). The devices were evaluated using an agar phantom and in hindlimbs of anesthetized rabbits. This brief report presents the use of these proof-of-concept semi-implantable devices in the limbs of healthy participants, a first-in-human demonstration of devices wirelessly powered and operated by volume conduction.

## II. Methods

### 1. Electronic system

The electronic system consists of 1) an external system that delivers the HF current bursts for WPT and bidirectional communications, and 2) wireless devices for EMG sensing and electrical stimulation. This electronic system was extensively described in (17). Yet some aspects are highlighted here to facilitate the reading of this first-in-human demonstration.

#### External system

The selected hierarchical architecture for the BHNS proposes the use of an external system that consists of one top-level controller that communicates with the wireless AIMDs through external low-level control units. These control units apply HF current bursts to the tissues to power and communicate with the wireless devices. The currents are applied using external textile electrodes strapped around the limbs (Fig. 1a).

The implemented control unit delivers HF bursts for powering. They include a single 30 ms “Power up” burst to power up the wireless implantable devices located between the two external electrodes, and power maintenance bursts with a repetition frequency (*F*) of 50 Hz and duration (*B*) of 1.6 ms to keep them energized. To perform bidirectional communications, a custom communication protocol stack was created. In the case of downlink (i.e., information sent from the external system to the wireless devices), the HF bursts are amplitude modulated by the external system. A set of 13 different commands can be used to 1) ask if an implant with a specific address is active (hereinafter referred to as “Ping”), 2) configure the sensing and stimulation parameters, 3) control their execution, and 4) uplink EMG samples. In the case of uplink (i.e., information sent from a wireless device to the external system), the external system delivers HF currents that are modulated – by means of load modulation – by the wireless devices to generate an uplink reply. This is seen by the control unit of the external system as minute changes in current.

The duration of the downlink and uplink bursts, as well as the EMG sensing and stimulation parameters that can be set from the external system are reported in (17). The shortest uplink message is that replied after the command “Ping”. During downlink, the external system sends this command to a specific device identifier. The implants located between the two external electrodes receive this information, and if their address matches that sent by the external system, an 8-bit frame acknowledge (ACK) is uploaded. In the case of the downlink “Get Config” command, which is used to know the current EMG acquisition or stimulation configuration of the device, the floating device replies with an uplink message encoded in 3 bytes. The longest uplink message is received when the external system requests an EMG sample. The addressed device replies with 4 bytes. Because of this, “Ping” is the initial command used to test the performance of bidirectional communications. If the performance is adequate, more complex commands (i.e., 3 and 4 bytes) are tested.

#### Wireless devices for EMG sensing and electrical stimulation

As explained in (17), the proof-of-concept wireless device for the acute human demonstration reported here consists of ultrathin intramuscular electrodes based on thin-film technology. Their biocompatibility was previously demonstrated in cytotoxicity, sensitization and irritation tests according to ISO 10993 (23). The intramuscular electrodes are connected to a miniature electronic circuit to be fixed on the skin. Each device has only two double-sided electrodes on a narrow polyimide filament (coined Electrodes ‘A’ and ‘B’ in Fig. 1c) for powering, bidirectional communications, electrical stimulation, and EMG sensing. This main filament has a width of 0.42 mm, a length of 81.6 mm, and a thickness of 0.02 mm. Each electrode contact has a width of 0.265 mm and a length of 7.5 mm, and their edges are rounded to avoid sharp corners that would lead to high current densities. The distal contacts located in the top and bottom layers of the main filament are electrically connected in the miniature electronic circuit to use them as a single distal electrode (Electrode ‘A’), with a total surface area of 3.8 mm^2^. This is also done with the two proximal contacts located in the top and bottom layers of the filament, creating a proximal electrode of 3.8 mm^2^ (Electrode ‘B’). The impedance magnitude of the electrodes, as measured across two electrode pairs in 0.9% NaCl, is flat beyond 10 kHz, and lower than 190 Ω at 1 MHz (17). The distance between the electrodes’ centers is 30 mm. To facilitate the insertion of the electrodes in the muscle, the distal end of the electrodes’ filament has a U-shape structure that ends in a guiding filament (Fig. 1c). This guiding filament is inserted through the lumen of a 23 G hypodermic needle (Sterican 4665600 by B. Braun Melsungen AG) having a length of 60 mm and an outer diameter of 0.6 mm, and the main filament with the active electrodes runs externally to the needle. As the guiding filament is so thin and the U-shape lies very close to the needle’s bevel, the bevel was smoothed with a laser (Picco Laser, by O.R. Lasertechnologie, Germany) prior to inserting the guiding filament inside the needle and towards the needle’s hub.

The miniature electronic circuit connected to the intramuscular electrodes (Fig. 1b) includes a microcontroller that has been programmed to run in three power consumption modes to guarantee energy efficiency. The current consumption measured during idle mode, processing and basic operations mode, and sensing mode are 180 μA, 185 μA, and 400 μA respectively (17). With this current consumption, it is possible to power all the electronic components and the control unit’s peripherals required for the power mode defined. At first, the device is configured to power up and wait for instructions sent from the external system (i.e., idle mode). The power up is done using the 30 ms “Power up” burst, and the device is kept energized with the power maintenance bursts (burst frequency *F* of 50 Hz, and duration *B* of 1.6 ms) using a set of two 10 μF capacitors connected in parallel, which provides a stable dc voltage for the control unit and the rest of the electronics during and in-between HF bursts. To perform electrical stimulation or EMG sensing, the device must be first configured from the external system. In the case of electrical stimulation, the external system can program the device to deliver biphasic or monophasic pulses, and the direction of the pulses. The external system sets the frequency of stimulation, the pulse width of the stimulation pulse, and the interphase dwell in case a biphasic symmetric pulse is required. This is done by modifying the frequency and duration of the applied HF bursts during the stimulation phase. Usually, the users set the stimulation frequency between 30 and 200 Hz, the pulse width between 100 and 200 µs, interphase dwell of 30 µs, and cathodic-first pulse. The current limiters inside the electronic circuit fix a maximum stimulation amplitude of 2 mA. In the case of EMG sensing, the external system must configure the type of EMG acquisition to be done (raw, or 2 options of parametric acquisition), the sampling frequency (250, 500, 750 and 1000 Hz, similar to those reported in (24–27)), and the window size in case the parametric option is set (15 different options starting at 10 ms, up to 500 ms, similar to those reported in (1,28–30)). These specific parameters (i.e., algorithm for parametric acquisition, possible sampling frequencies and windows) can be changed by reprogramming the control unit of the proof-of-concept device using the 6 programming pads accessible from the top layer of the circuit. When the EMG acquisition is started from the external system, the wireless device acquires a sample and stores it in the random-access memory (RAM) of the control unit (ultra-low power microcontroller, MKL03Z-32CAF4R by NXP Semiconductors N.V.). Because of the size of the microcontroller, this memory is limited to 2 kB. When acquiring in parametric mode, the parametric value is calculated, and the result obtained for a sampling window is saved in memory, improving the efficiency of memory usage.

The analog front-end (AFE) designed is explained in detail in (17). It has been configured to have a gain of 54 dB, a signal-to-noise ratio (SNR) of 46 dB, and a common mode rejection ratio (CMRR) of 88 dB. The circuit board includes a test point to access the output of the AFE for debugging. The AFE’s output is connected to the control unit’s analog-to-digital converter (ADC), which is programmed to have a resolution of 10 bits. This resolution can be increased to 12 bits by reprogramming the device with the programming pins. During EMG sensing, the control unit monitors the output of its downlink demodulator to identify if a HF burst is being applied by the external system. If this is the case, the AFE’s output is saturated by the HF bursts for a very short period of time (1.6 ms, equivalent to the duration of the burst), and then recovers from it in approximately 4 ms. When this saturation artifact happens, the control unit replaces the samples corresponding to this saturation with a constant value (e.g., five samples for a sampling frequency of 1 ksps), without affecting the sampling frequency defined by the external system. By doing this blanking (31), it is possible to identify saturation and recovery time windows without misinterpreting this effect with high amplitude EMG activity.

Fig. 2 shows the different sequences that can be obtained with the interaction of the external system and its low-level controller, and the wireless implants that are within the textile electrodes of this controller. During downlink, the information is modulated in the HF currents, and is demodulated by the implant (Fig. 2a). If an ACK reply is requested (e.g., with the “Ping” command), the external system delivers a burst, and the implant does load modulation to uplink this information, which is seen by the control unit as minute changes in current consumption (Fig. 2b). For stimulation, the external system delivers HF bursts with the pulse width and repetition frequency defined for stimulation, and the implant rectifies these volume-conducted HF bursts to cause local low frequency currents capable of stimulating excitable tissues. Fig. 2c shows a low-pass filtered biphasic stimulation waveform obtained *in vitro* using an oscilloscope (17). For EMG acquisition, the external system delivers power maintenance bursts (50 Hz, 1.6 ms) which causes a short saturation in the AFE of the implant, followed by a time window for EMG acquisition (Fig. 2d). The example shows the output of the AFE (i.e., AFE test point) obtained *in vitro* using an oscilloscope when no EMG signal is present across the intramuscular electrodes (17). The recorded samples include the samples with saturation artifacts (samples to be blanked by software) and the valid samples obtained during the acquisition window.

**Fig. 2.**
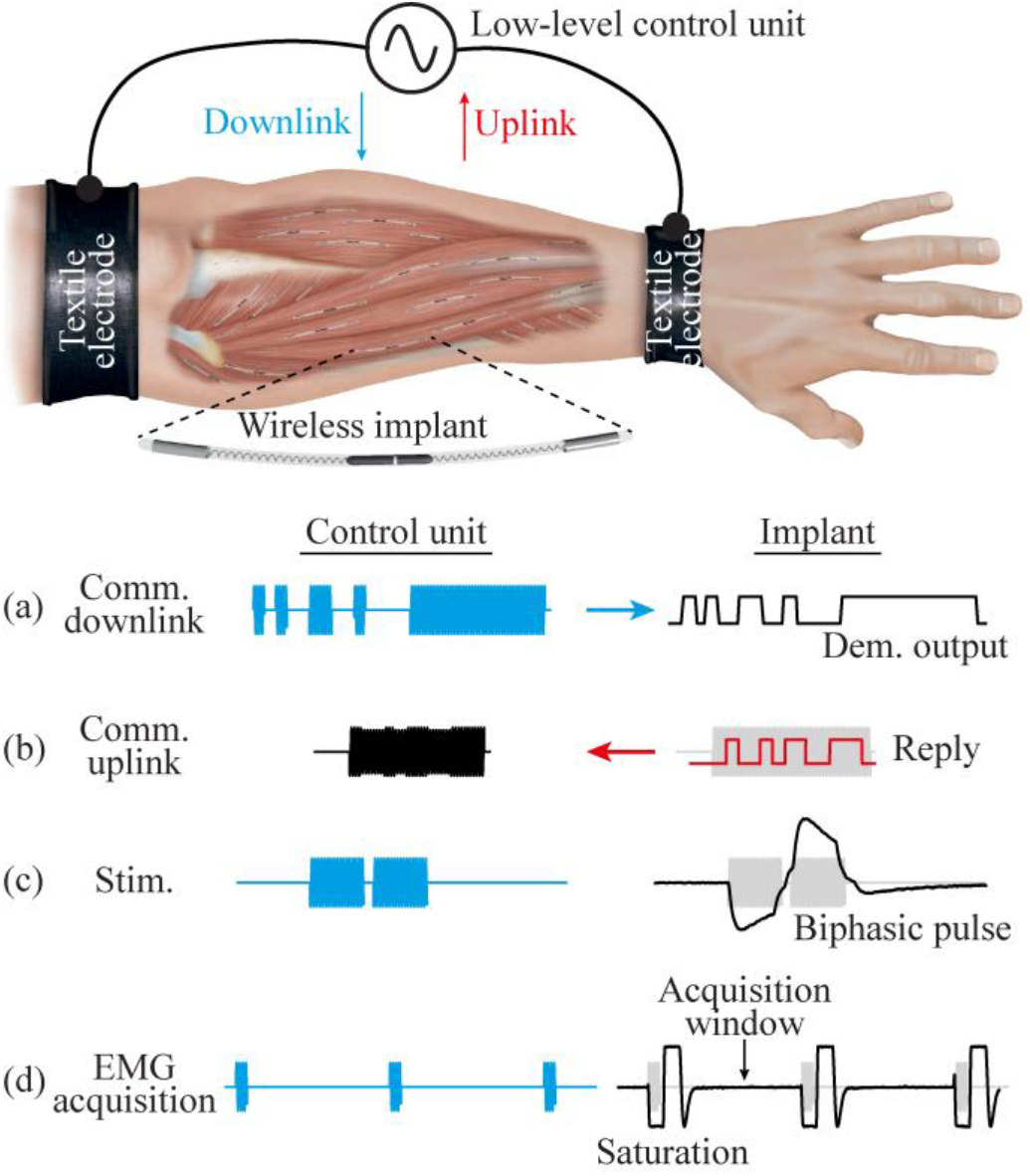
HF current bursts sequence. It shows the different sequences that can be obtained with the combination for the external system and its low-level control unit, and the wireless implantable devices. See main text for detailed information.

For this first-in-human demonstration, the proof-of-concept devices were double blister-packaged in clear Stericlin bags and heat sealed. The devices were sterilized by Ethylene oxide gas sterilization (8%) at low temperatures (38 – 42 °C).

### 2. Participants

The assays were performed at the Movement Disorders Clinic of Gregorio Marañón Hospital (Madrid, Spain) for the upper limb and at the Biomechanics and Assistive Technology Unit of National Hospital for Paraplegics (Toledo, Spain) for the lower limb. The procedures were conducted in accordance with the Declaration of Helsinki and approved by the local Ethics Committees (reference numbers 18/2020 and 565 respectively), as well as by the Spanish Agency of Medicines and Medical Devices (AEMPS) – records 858/20/EC and 856/20/EC respectively. Three healthy volunteers participated in the upper limb assays (23, 31, and 47 years old, all male), while other three healthy volunteers participated in the lower limb assays (29, 32 and 37 years old; two males, one female). The volunteers were recruited through a call for participation sent by email to colleagues and by another call posted in the facilities of both hospitals, and were not paid for their participation in the study. Before starting the experimental procedure, the participants were provided with oral and written information regarding the study (including risks, contingency measures, benefits, and data protection aspects) and signed an informed consent form.

### 3. Experimental Procedure

At first, the target muscle was defined for each participant. In the case of the upper limb, the target muscle was the biceps brachii or the triceps brachii; while in the lower limb it was the tibialis anterior, the gastrocnemius medialis or the gastrocnemius lateralis. A single semi-implantable floating EMG sensor and stimulator was set up per participant. The electrodes were implanted using the dedicated 23 G hypothermic needle (Fig. 1c). The approximate site for the deployment was defined using anatomical cues. The criteria were 1) that the electrodes were aligned to the applied electric field, 2) that at least 40 mm of the needle were completely inside the muscle to guarantee that both electrodes were in this tissue, and 3) if possible, to insert the tip of the needle – which is few mm away from Electrode ‘A’ – close to the motor point of the target muscle. Before inserting the thin-film electrodes of the semi-implantable device, the area was cleaned with chlorhexidine. This procedure was guided by ultrasound to verify that the needle was correctly inserted in the muscle belly, and that its orientation was as parallel as possible to the skin (to maximize the alignment of the electrodes with the applied electric field delivered using the external textile electrodes). After the needle was inserted in the target location, the miniature electronic circuit (enclosed in a polymer capsule for protection) was fixed to the skin of the participant using adhesive tape to avoid the accidental extraction of the electrodes. Finally, the end of the guiding filament adhered to the needle hub was cut, and the needle was gently extracted. Fig. 3 shows proof-of-concept semi-implantable devices deployed in the biceps brachii of participant 2, and in the gastrocnemius medialis of participant 5.

**Fig. 3.**
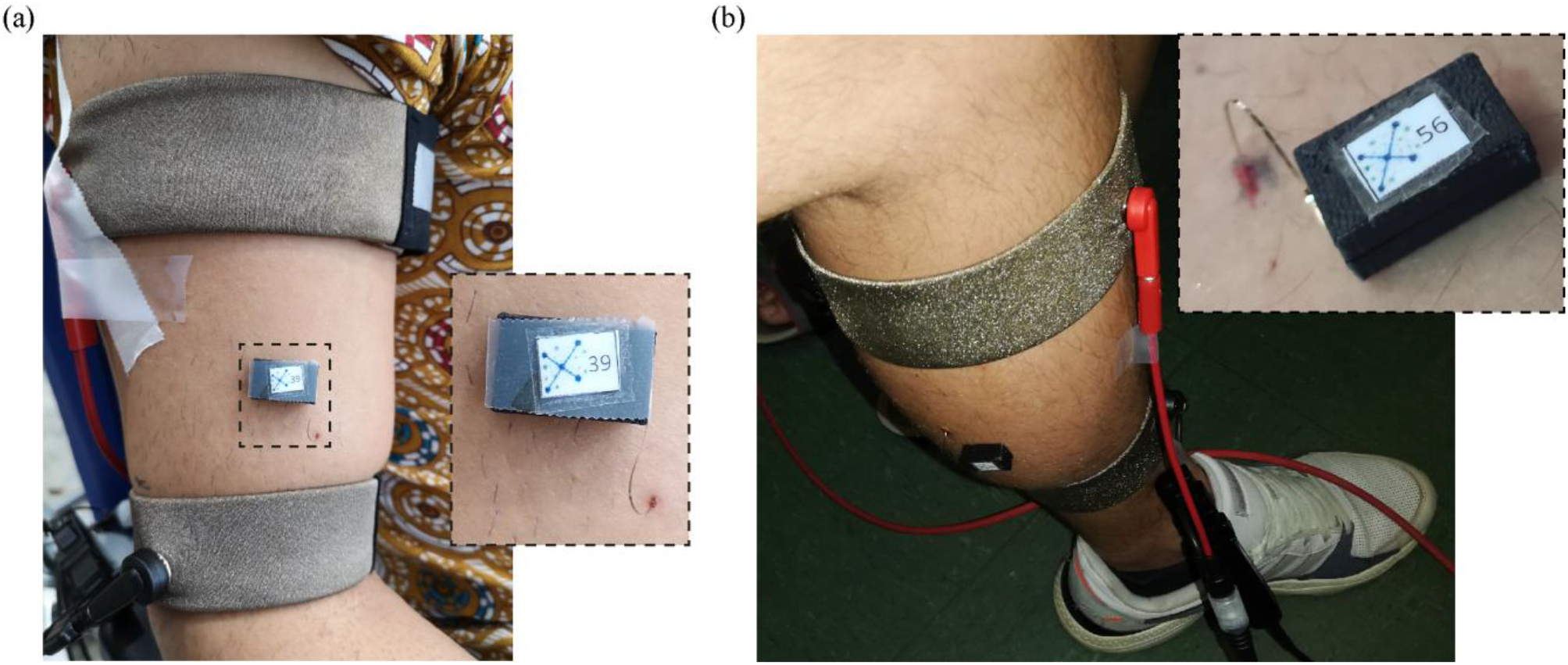
Proof-of-concept semi-implantable devices deployed in the target muscles of two participants. a) Device in biceps brachii of participant 2. b) Device in gastrocnemius medialis of participant 5.

Once the intramuscular electrodes were successfully inserted in the muscle belly, the participants remained seated on a chair. Then, the textile electrodes were strapped around the limb and were connected to the external system. In the case of the upper limb assays, the participant was asked to keep the upper arm aligned with the torso, to lay down the forearm on the armrest (thus forming a 120° angle in the elbow), and to keep the wrist in neutral position.

Several tests were performed to test the ability of the external system to power and bidirectionally communicate with the proof-of-concept device. To evaluate the bidirectional communications, it has been defined the success rate: the number of messages correctly decoded by the external system (testing the downlink and uplink sequence), divided by the number of messages that should have been theoretically received. If any of the error detection mechanisms available in the external unit is triggered (i.e., error in Manchester coding, parity bit, frame length or command code), the message is marked as incorrect. In the case of a “Ping”, the success rate would only evaluate the 8-bit frame ACK. In much more complex commands and subsequent replies such as “Get Config”, the message is understood as the total amount of information (3 bytes or 4 bytes) to be received by the external system.

At first, the external system was set at an initial peak amplitude of 30 V, and it was configured to deliver the “Power up” burst and “Pings” requests. If the address sent during downlink matched the address defined in the semi-implantable device, this device sent an ACK frame to the external system. The peak amplitude was then increased until uplink waveforms with the ACK frame were seen in the demodulator’s shunt resistor of the external system (Fig. 4a). If this bidirectional communication trial was successful (i.e., success rate above 90%), longer communication frames were tested to guarantee that the external system could correctly configure the semi-implantable device for electrical stimulation and EMG sensing, trigger these functions (i.e., stimulate, start/stop sensing) and upload the samples of EMG activity acquired by the floating device.

**Fig. 4.**
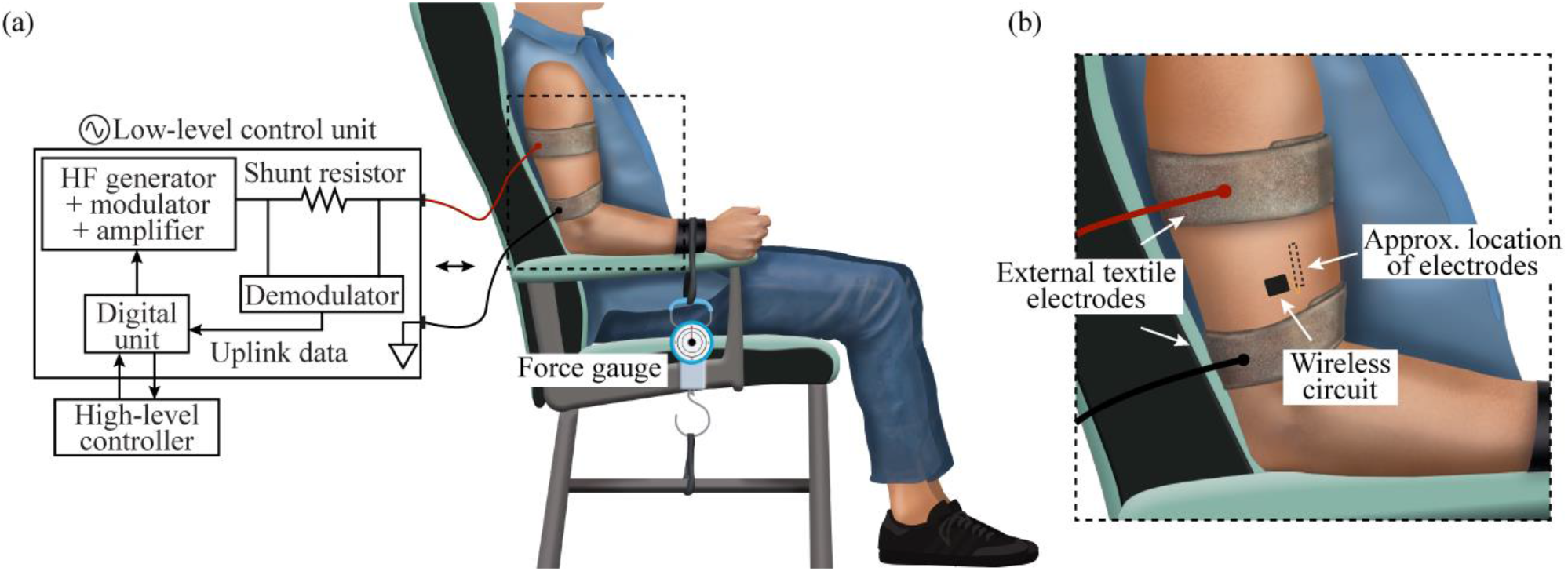
Setup used during the study in upper limb. The participant was always sitting on a chair, with the upper arm aligned with the torso and the elbow at an angle of 120°. The strapped external textile electrodes were connected to the external system. a) For the force measurement during isometric contractions of the biceps brachii, the wrist was strapped to the armrest of the chair, and to a force gauge. b) Zoom of the region where the wireless circuit was located, and approximate location of thin-film electrodes.

After it was confirmed that the semi-implantable device was properly powered by the external system, and that the bidirectional communications were successful, electrical stimulation and EMG sensing assays were performed. In the case of electrical stimulation assays, videos were recorded to capture the moment in which a stimulation sequence was triggered from the external system. In the case of EMG sensing for biceps brachii assays, it was not possible to assess muscle contraction using surface electrodes connected to a commercial EMG amplifier as the amplifier saturated with the HF current bursts applied – as expected – but it could not return from the saturation fast enough to acquire EMG signals in-between bursts. For this reason, a force gauge was used to measure the tensile force exerted by the arm when the biceps brachii was contracted with the hand closed. The force gauge was attached firmly to the participant’s arm – which was held in the armrest with an atraumatic band – and its base was fixed to a base of the chair, forcing an isometric contraction (Fig. 4). The participant was requested to perform different force profiles, such as maximum sustained contraction, target forces (e.g., 3 kg, 5 kg, 10 kg), and sequences of hard and short contractions. In the case of target forces, the obtained measurements were read aloud as a means of feedback for the participant. During the contractions, EMG was acquired by the floating device, and the samples were uploaded to the external system for offline analysis.

Each participant was constantly asked if he or she perceived any type of sensation related to the HF current bursts applied or to the floating device implanted, and the region close to the textile electrodes was visually monitored continuously. The surveillance also included measurements of the impedance across the external textile electrodes to identify possible changes in the interface between the textile electrodes and the skin. This was done using an active differential probe (TA043 from Pico Technology Ltd) to measure the voltage applied across the external electrodes, and a current probe (TCP2020A from Tektronix, Inc.) to measure the HF current flowing through the tissues. Both probes were connected to a battery-powered oscilloscope (TPS2014 from Tektronix, Inc.). After finalizing the assays, the external system was turned off and the textile electrodes were unstrapped. Then, the intramuscular electrodes were extracted by holding the guiding filament and the electrodes’ filament and gently pulling out, and the insertion region on the skin was cleaned with chlorhexidine. The participants remained for 30 more minutes in the hospital for monitoring, and were monitored again 24 h later through a phone call to ensure their well-being and to note possible adverse effects.

## III. Results

### 1. Implantation

The insertion mechanism was minimally invasive and did not require local anesthesia. In the case of the implantation procedure in the triceps brachii, due to limitations in the room, the position during implantation was uncomfortable for the physician. This hampered the correct insertion of the electrode, which broke during the implantation. In the case of the tibialis anterior and gastrocnemius lateralis muscles, the intramuscular electrodes broke during insertion as the dedicated needle experienced more stiffness due to the insertion angle required to maximize the alignment of the electrodes with the applied electric field. For these reasons, the study was finally performed only in the biceps brachii and the gastrocnemius medialis. In the cases in which the implantation process failed, the broken thin-film electrodes and the needle were gently extracted by simultaneously pulling from the main filament, the guiding filament and the needle hub. Table 1 summarizes the final location of the proof-of-concept device in all the participants.

**Table 1.** Demographic information and bidirectional communications success rate in each participant.

| Participant | Gender | Age | Target muscle | Communications success rate |
| --- | --- | --- | --- | --- |
| 1 | Male | 47 | Biceps brachii | No communication |
| 2 | Male | 31 | Biceps brachii | Ping: 100%<br>Get Config: 100% |
| 3 | Male | 23 | Biceps brachii | Ping: 100% |
| 4 | Female | 37 | Gastrocnemius medialis | No communication |
| 5 | Male | 29 | Gastrocnemius medialis | Ping: 40% |
| 6 | Male | 32 | Gastrocnemius medialis | Ping: 80% |

Fig. 5 shows the ultrasound image obtained immediately after the needle was placed in the biceps brachii of participant 2. Only 3.4 cm of the 6 cm needle are seen in the image (limited by the ultrasound’s transducer length), but the resulting insertion length was approximately 5.5 cm. The participant had a subcutaneous fat layer of approximately 3 mm, and the needle – corresponding to the position of the thin-film electrodes – had an angle of approximately 8° with respect to the skin.

**Fig. 5.**
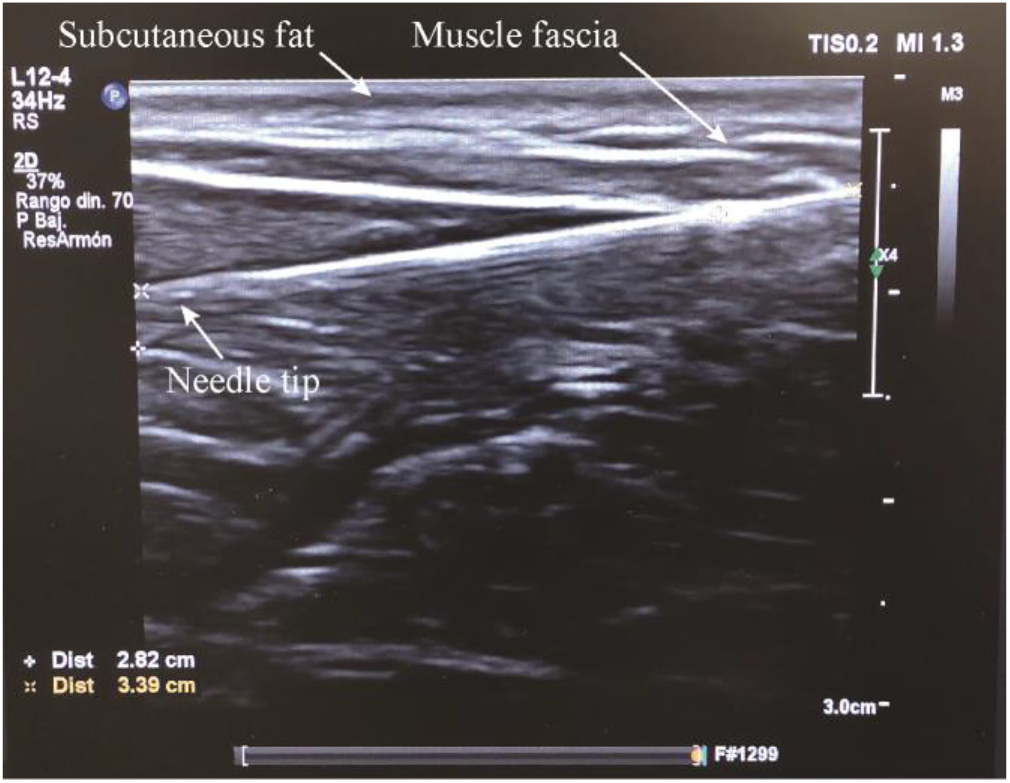
Ultrasounds image obtained from participant 2 immediately after the needle was inserted in the biceps brachii. The needle was injected from the top right corner (not shown in the image), passed through subcutaneous fat and the muscle fascia, and ended up in the target muscle. The tip of the needle is seen at the left of the image at a depth of approximately 1 cm.

After the implantation, the participants did not report discomfort due to the intramuscular electrodes. The filament was thin enough so that the participants could move the limb under study without any type of sensation. For this same reason, the participants were asked to remain seated until the study was finished to avoid accidental extraction.

### 2. Bidirectional communications

Fig. 6 shows an example of the waveforms obtained from the external system when it requested a “Ping”, and the floating device replied with an ACK frame. The waveforms were obtained in the upper arm (Fig. 6a) and the lower leg (Fig. 6b and Fig. 6c) of three different participants. The external system was delivering HF bursts at a frequency of 1.1 MHz (power maintenance burst frequency *F*: 50 Hz, and duration *B*: 1.6 ms). For any uplink, the external system delivers a HF current burst to trigger the reply and give energy to the semi-implantable device (Fig. **6**, blue waveform). The load modulation made by the floating device creates minute changes in the current flowing through the external textile electrodes. This current is measured by the external system’s demodulator using a shunt resistor (Fig. 4). Fig. 6 – orange waveform – shows the output of the demodulation circuit after filtering and amplifying the voltage across this shunt resistor. This output later passes by an analog comparator, whose output connects to the receiver of the external system’s universal asynchronous receiver-transmitter (UART) to decode the information sent by the semi-implantable device. In the case of participant 2 (Fig. 6a), who had the semi-implantable device deployed in the biceps brachii, success rates of 100% were obtained for bidirectional communications, both in short frames as the “Ping” command, as well as in longer frames such as “Get Config” when the external system was delivering bursts with a peak amplitude of 60 V and 1.42 A, and the textile electrodes were separated 10 cm. It is worth noting that these peak amplitudes for electric potential and current are scaled down due to the use of bursts that have a duty cycle of 0.08 (power maintenance burst frequency *F*: 50 Hz, and duration *B*: 1.6 ms). For another upper limb participant, participant 3, success rates of 100% were obtained for pings to a device implanted in the biceps brachii. In the case of participant 5 (Fig. 6b), whose floating device was in the gastrocnemius medialis, the maximum success rate for pings was only 40% (68 V, 2.28 A, 10 cm distance between centers of textile electrodes). Its orange waveform clearly shows a HF component in the uplink waveform (i.e., output of demodulator of external system), which may have generated an incorrect digital signal in the analog comparator of the external system’s control unit, therefore generating errors in the decoder. Additionally, the waveform shows low peak-to-peak variations during the ACK, which hampers even more the decoding of the data. In the case of participant 6 (Fig. **6**c), whose device was also implanted in the gastrocnemius medialis, the maximum success rate for pings was 80% (90 V, 2.1 A, 12 cm distance between textile electrodes). The orange uplink waveform also contains some HF component, but the ACK ping message differs clearly from the rest of the window.

**Fig. 6.**
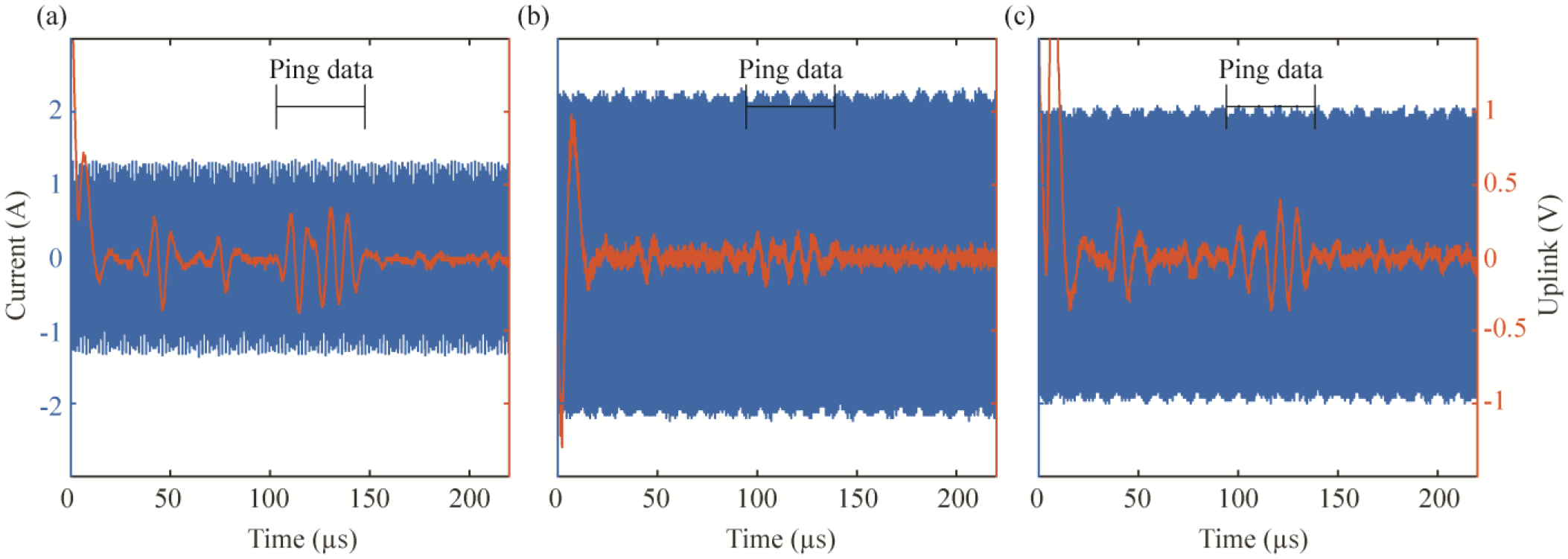
Waveforms obtained with a floating oscilloscope and the probes connected to the external system. These waveforms are obtained after the external system requests a “Ping” to a specific device. In blue, HF current delivered by external system for uplink; in orange, output of external system’s demodulator, showing the filtered and amplified voltage across the shunt resistor when the floating device does an ACK ping message by load modulating the HF current burst (i.e., shows changes in the current flowing through the textile electrodes). a) Participant 2, biceps brachii. b) Participant 5, gastrocnemius medialis. c) Participant 6, gastrocnemius medialis.

The “Ping” requests sent to the device implanted in the biceps brachii of participant 1 and the gastrocnemius medialis of participant 4 were not properly received by the external system. In both cases it is unclear whether the devices were not properly powered (e.g., because the proximal electrode (‘B’ in Fig. 1c) was lying in fat tissue), or whether the hardware-based demodulation and decoding circuit of the external system was not able to properly acquire and interpret the signal modulated by the floating devices when they were sending their ACK ping message (e.g., because the modulation index was too low for the digital converter to work properly).

### 3. Electrical stimulation and EMG sensing

As the semi-implantable device in participant 2 was able to be powered by and to bidirectionally communicate with the external system in both short and long communication frames, it was possible to configure the stimulation and EMG acquisition parameters of the floating device. A video was recorded during a stimulation sequence when the floating device was configured to do biphasic symmetric stimulation with a cathodic-first pulse (i.e., a cathodal phase is followed by an anodal phase to obtain a charge-balanced pulse (32)). This sequence is shown in the schematic representation of the stimulation shown in Fig. 2c. The external system delivered HF current bursts to generate a stimulation with a frequency of 150 Hz, 120 µs pulse width and interphase dwell of 30 µs, with a maximum stimulation amplitude of 2 mA. When the stimulation was triggered by the external system, a visual feedback confirmed that the skin area over the region where the thin-film electrodes were located slightly sank, indicating muscle contraction. Analyzing the video frames, it is estimated that the skin area sank approximately 1.2 mm during stimulation.

In the case of EMG acquisition, it was possible to obtain raw EMG. Fig. 7a shows two raw waveforms recorded by the wireless device and transmitted to the external system: the baseline (blue) and a voluntary maximum sustained contraction done by the participant (orange). The floating device was configured to acquire raw EMG at a sampling rate of 1 ksps. The method for blanking the artifacts during the AFE’s saturation and recovery allowed us to distinguish this time window and correctly obtain EMG recordings during the acquisition window (Fig. 2d) despite the effect of the HF current burst in the output of the AFE.

**Fig. 7.**
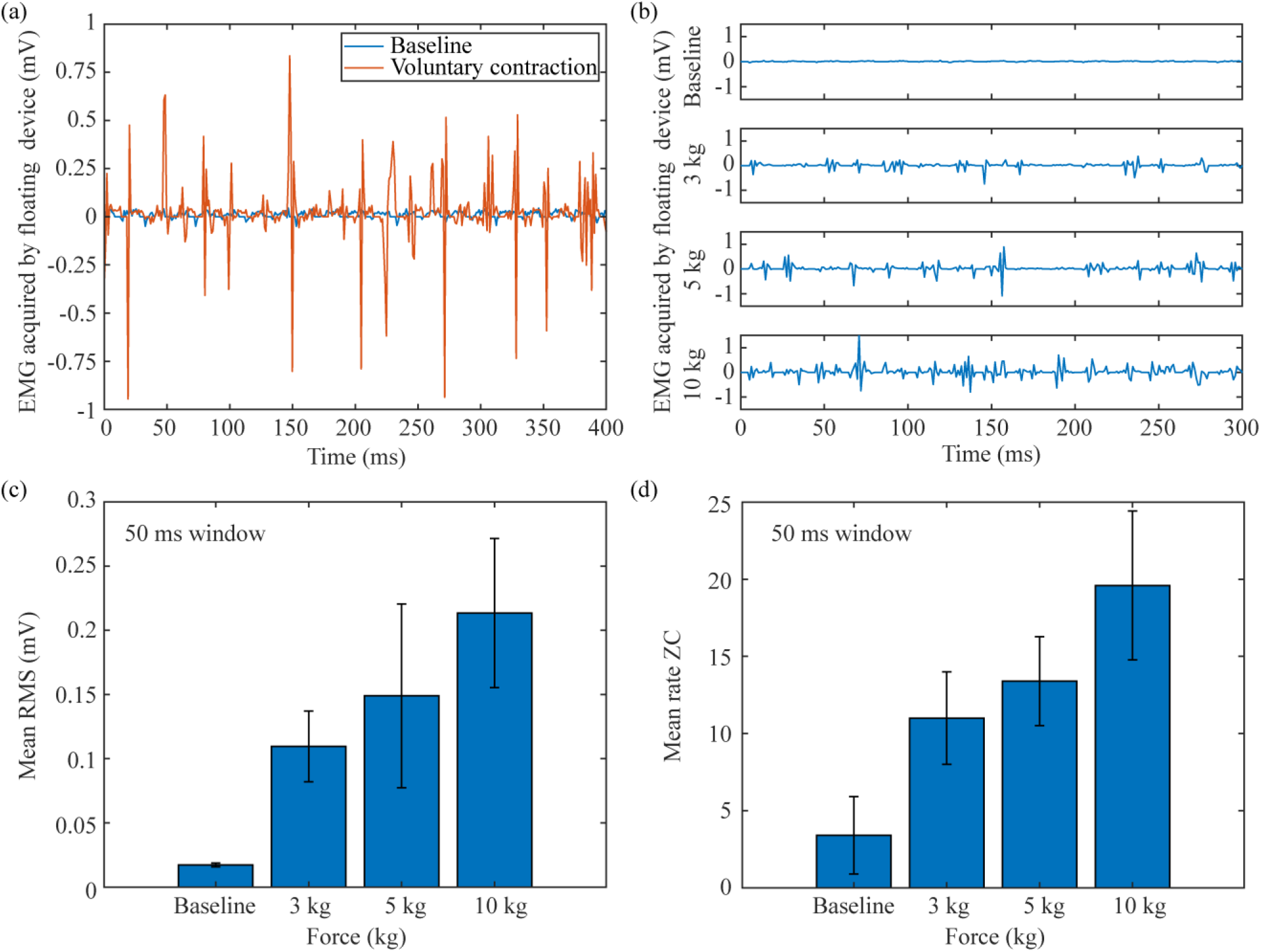
Examples of raw EMG obtained by the floating EMG sensor and stimulator in biceps brachii of participant 2. A) Comparison between baseline and sustained contraction. b) Comparison between baseline and three different target forces exerted by the participant. c) Offline calculation of average of root mean square (RMS) value for the raw signals shown in b. d) Offline averaging of zero crossing (ZC) calculation for the raw signals shown in b. In both cases, the window used for calculation was set to 50 ms.

Using a force gauge as a reference (Fig. 4a), participant 2 was asked to contract his arm to target a specific force. Fig. **7**b shows raw EMG recordings obtained for four trials, corresponding to no contraction (baseline), 3 kg, 5 kg and 10 kg of force. As the target force increases, the electrical activity measured from the floating device increases. These raw signals (Fig. 7b) were used to quantify offline the root mean square (RMS) value and the zero crossings (ZC) rate to evaluate its use in a future BHNS. RMS is an amplitude estimator while ZC, calculated by finding the rate of crosses by zero in the time domain, is an indicator of the changes of the spectral content of the EMG (33). The recorded signals were divided into 50 ms windows, and for each window the RMS and ZC values were calculated. Fig. **7**c-d show the mean RMS and mean ZC respectively obtained for each raw signal. The threshold for the offline ZC algorithm was defined as 0. As expected, both parameters increased when the force increased.

To avoid the limitations imposed by the internal memory of the miniature electronic circuit of the floating device, and to be more efficient with the communication channel by minimizing the amount of data to upload for the control of the BHNS, the floating device could internally calculate a parametric value of the obtained EMG. To limit the calculation time of the control unit (i.e., the amount of time it must be active in its higher power mode), it was considered that the parameters should be obtained from the signal in the time domain. One of these parameters is the rate of ZC, which is easier to implement in hardware (34). A trial of parametric EMG based on ZC was done to evaluate the possibility of performing longer recordings. The floating device was configured to do EMG acquisition at a sampling rate of 1 ksps and parametrize the results using ZC with a window size of 30 ms, a window size that could be used for real-time control. The participant was asked to do three hard and short contractions, followed by relaxations, in cycles of 2 s in a 6 s frame (progressive contractions starting approximately at seconds 0, 2 and 4), while the semi-implantable device was acquiring the information. Fig. 8 shows the result of this assay, in which the three progressive contractions can be clearly noticed. An offline moving average filter of 15 samples was used to highlight the envelope of the obtained parametric EMG.

**Fig. 8.**
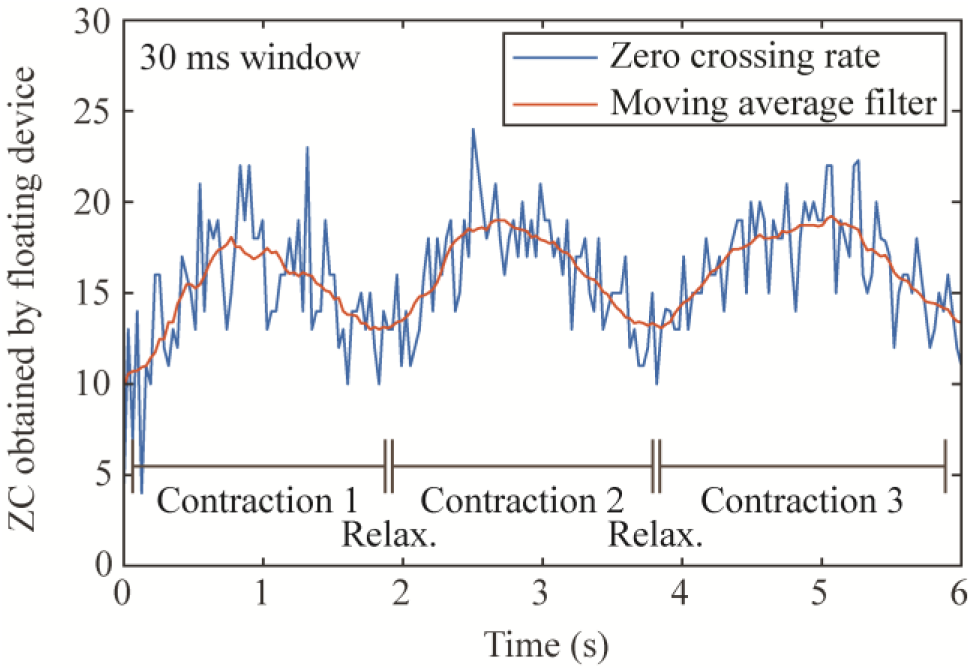
Parametric acquisition using ZC rate. Participant 2 did three contractions during 6 seconds (progressive contraction followed by relaxation), which can be clearly seen by the parametric waveform obtained. After the samples were uploaded to the external system, a moving average filter was added to show the envelope of the contractions.

During the trials to assess bidirectional communications, electrical stimulation and EMG acquisition, the participants did not report any type of perception related to the application of the HF current bursts. This aligns with the results obtained in (21).

### 4. Extraction

After the assays were performed on the upper limb or lower limb of the participants, the intramuscular thin-film electrodes were gently extracted. The electrodes sled smoothly through the tissues, and both the main filament with the active sites of the electrodes and the thin guiding filament were extracted completely. There was no bleeding, pain, or secondary effects.

## IV. Discussion

Here it is presented a brief report of the first-in-human demonstration of floating devices powered and controlled by volume conduction, which are capable of EMG sensing and electrical stimulation. These proof-of-concept semi-implantable devices were designed for the acute human study presented here and their technical implementation and *in vivo* validation were previously described (17).

Frequent breakage of the thin-film electrodes during their implantation was one of the most important limitations of the present study. In contrast to previous studies (upper limb (1,35), lower limb (36–38), neck (36,39), and hand (36,38) muscles) in which very similar thin-film electrodes were reproducibly implanted transversally to and/or shallower into the muscle, here the electrodes had to be deeply inserted longitudinally to the limb to guarantee 1) that both the distal and proximal electrodes (‘A’ and ‘B’ respectively in Fig. 1c) were inside muscle tissue, and 2) that the power picked up by the floating device during the assays was enough to power the miniature electronic circuit. To accomplish this, the needle had to be completely inserted as longitudinal as possible to the muscle, in acute angle with the skin and the muscle fascia. Remarkably, the muscle fascia is an anisotropic connective tissue (40–42) that has higher stiffness and strain in the longitudinal than the transverse direction (43). This implies that the needle and the thin guiding filament may have faced higher mechanical resistance in this insertion scenario than in the typical scenario of use for this kind of thin-film electrodes (i.e., perpendicular to the muscle). In the case of the lower limbs the conditions are more detrimental, as the fascia (e.g., crural fascia in the lower leg) is thicker and stiffer because it helps stabilize the limb during locomotion (44), and lower limb muscles are stiffer than upper limb muscles, making it even more difficult to insert the dedicated needle. It is worth noting that the crural fascia in the anterior compartment (tibialis anterior muscle) is stiffer than in the posterior compartment (gastrocnemius medialis muscle) (45). This could explain why it was more difficult to implant in the tibialis anterior than in the gastrocnemius muscles. It has been also experimentally determined that the stiffness of the gastrocnemius lateralis muscle is higher than that of gastrocnemius medialis muscle, which could explain why it was easier to implant the device in the gastrocnemius medialis muscle (46,47).

Even though this thin-film technology has been implanted in animals for recording purposes for up to 9 weeks (48), and in humans for electrical stimulation studies for up to 11 weeks (49), it must be noted that the devices presented here are proof-of-concept prototypes that demonstrate the circuit architecture and the capabilities of the proposed approach in an acute study. They are not the final proposed implantable devices to be chronically implanted. The electronics are meant to be integrated in the future into an ASIC to be included in a flexible threadlike implant with two electrodes at opposite ends as that shown in (22). In other words, the limitations faced with the thin-film electrodes will not be faced with a conformation based on a tubular body mostly made of silicone.

In this first-in-human study a single floating device was used per participant. This limitation was mainly due to timing restrictions in the hospitals where the study was performed. It was considered sufficient to inject only one wireless device for EMG sensing and electrical stimulation to demonstrate the WPT and communication method based on volume conduction. However, it must be noted that the designed communication protocol allows the control of up to 256 devices per low-level control unit, and the possibility to have several low-level units connected to a high-level controller. In fact, we have *in vitro* demonstrated that more than 10 floating devices can be wirelessly powered and digitally controlled from the external system (50).

In terms of the results obtained for bidirectional communications, it is worth noting that the demodulation performed by the external system consists in a rather simple hardware-based approach: the output of the analog amplitude demodulator is interfaced with the UART of the digital unit using an analog comparator. This approach has several limitations, including fixed thresholds and the impossibility to postprocess the digital data to recover lost bits. The limitations hinder the control unit’s decoding performance in certain scenarios, which translates into very low success rates. For example, as the electric potential across the thin-film electrodes decreases, the modulation index seen from the external system during uplink decreases too. This can be improved by replacing the current approach used in the demodulator of the external system (i.e., analog comparator plus UART) with a more flexible approach based on digital processing: digital filters, comparators with adaptive thresholds and specific decoding algorithms. This could increase the success rate for uplink in unfavorable cases where low SNR is obtained.

The muscle contractions obtained during electrical stimulation were weak compared to those obtained in previous electrical stimulation studies using this WPT approach (17,22). Because of the limitations explained above related to the frequent breakage of the thin-film electrodes when implanting the floating device, during the procedure it was prioritized the alignment with respect to the applied electric field and the settling in a single tissue over electrical stimulation (i.e., powering and bidirectional communications were deemed more important in the study). One limitation of using the same electrodes for powering, electrical stimulation and EMG sensing in the wireless device is that the HF current bursts applied by the external system saturate its AFE, creating an artifact during EMG sensing. This artifact is equivalent to that generated when electrical stimulation and EMG recording is done simultaneously in research and commercial systems (51). Several applications report this, including closed-loop myoelectric control using electrotactile stimulation (52), tremor suppression using EMG and electrical stimulation (53), and EMG acquisition during transcutaneous spinal cord electrical stimulation (54). Strategies to avoid stimulation artifacts include using sequential operations in which EMG recording windows are separated from electrical stimulation windows (1,53), and software blanking (31). The wireless device reported here uses a similar approach for the saturation artifacts: there are short windows in which the EMG samples acquired do not represent muscle activation as the HF current bursts generate a saturation artifact, and so, these samples are blanked replacing them with a constant and known value. This short, saturated window is followed by a long non-saturated EMG recording window, and the entire sequence is acquired at the sampling frequency previously defined from the external system.

The assays performed in the second participant were focused on transmitting raw EMG samples obtained by the floating device. However, it must be noted that raw EMG acquisition would not allow real-time control as the proof-of-concept floating device has a limited memory capacity of 2 kB, and all the samples must be uploaded to the external system, occupying the entire physical channel during several milliseconds (approximately 1.35 ms are required to upload one sample). This would limit the potential of the BHNS concept. It is worth noting that a similar uplink limitation is faced by other implants that do EMG acquisition (the IST-12 can upload at most 1 sample every millisecond (24)). The upload time could be decreased in the communication protocol by, for example, decreasing the duration of the power maintenance burst that is sent prior to the uplink transmission burst (more information in Additional file 1 of (17)). Also, by parametrizing the EMG signal in the floating devices it is possible to substantially reduce the number of bytes to be transmitted, thus allowing real-time control. We report an example using the rate of ZC, which is a basic parametric measurement obtained in the time domain. However, the communication protocol created allows us to use a second parametric measurement that can be implemented in the firmware of the floating devices or in an ASIC (e.g., RMS, spike properties (55)). Using parametric measurements, it is possible to acquire much more samples, store them in the wireless device’s memory, and be more efficient with the powering/communication channel to accomplish a BHNS system that can operate in real time to process and analyze the information obtained using the intramuscular electrodes, and activate the electrical stimulation or control external machines such as exoskeletons. By configuring the window size of these parametric measurements using the external system and its bidirectional communication protocol, the measurements can be adapted to different applications (e.g., selective and adaptive timely stimulation (SATS) for tremor reduction using RMS at a 10 ms or 15 ms window size (1,56), and autoencoding (AEN) of EMG for myocontrol using RMS at a 100 ms window size (57)).

The proof-of-concept floating AIMD described in depth in (17) and demonstrated here in the upper and lower limb of healthy participants cannot be used for chronic studies of more than 80 days, as the thin-film intramuscular electrodes may fail (58) and the skin access is prone to infections in the long term. However, the architecture of the miniature electronic circuit connected to these intramuscular electrodes can be integrated into an ASIC, as it is composed of typical electronic components that can be fabricated using current microelectronic technologies and manufacturing processes. This would allow to obtain fully implantable flexible threadlike EMG sensors and stimulators that can be easily implanted by injection, that can be employed in chronic studies and developed for future commercial use. We recently demonstrated in the hindlimbs of two anesthetized rabbits that fully injectable microstimulators (overall diameter of 0.97 mm) are able to rectify these volume conducted HF bursts to cause local low frequency currents capable of stimulation (22). The microstimulators include a hermetic capsule that houses an ASIC, two dc-blocking capacitors and a capacitance for voltage stabilization. The implantation procedure proposed, which uses a 14 G clinical catheter, is described in depth in (59).

## V. Conclusions

This brief report presents the first-in-human demonstration of the use of coupling by volume conduction as a WPT and bidirectional communications method to operate networks of wireless EMG sensors and stimulators. The proof-of-concept floating devices, consisting of thin-film injectable intramuscular electrodes connected to a miniature electronic circuit, were powered by and communicated with an external system that delivered HF current in the form of bursts via two external textile electrodes strapped around the limb where the device was deployed. In one participant out of six it was demonstrated that the external system can send digital commands to configure the EMG sensing and electrical stimulation parameters, to trigger the acquisition or the stimulation, and to upload samples. Strategies are proposed to improve the overall performance of the system, overcoming the limitations faced in the study regarding uplink speed and decodification, and memory usage. Even though the thin-film electrodes presented issues related to the need to insert them longitudinally to the limb, facing high mechanical resistance that led to breakage, it is worth noting that these limitations are inherent of these proof-of-concept prototypes. The demonstrated miniature electronic circuit will be integrated in an ASIC that will be included in a tubular implant with electrodes at opposite ends. The implantation procedure of this tubular conformation has been demonstrated to be successful in several assays, including those reported in (22). This first-in-human demonstration opens the possibility of using coupling by volume conduction in networked neuroprosthetic systems.

## List of abbreviations

AIMDs: active implantable medical devices
EMG: electromyography
HMI: human-machine interface
BHNS: Bidirectional Hyper-Connected Neural Systems
WPT: wireless power transfer
HF: high frequency
ASIC: application-specific integrated circuit
ACK: acknowledge
RAM: random-access memory
AFE: analog front-end
SNR: signal to noise ratio
CMRR: common mode rejection ratio
ADC: analog-to-digital converter
AEMPS: Spanish Agency of Medicines and Medical Devices
UART: universal asynchronous receiver-transmitter
RMS: root mean square
ZC: zero crossings;

## Declarations

### 1. Ethics approval and consent to participate

The procedures were conducted in accordance with the Declaration of Helsinki and approved by the local Ethics Committees of the Movement Disorders Clinic of Gregorio Marañón Hospital – Madrid – and the Biomechanics and Assistive Technology Unit of National Hospital for Paraplegics – Toledo – (reference numbers 18/2020 and 565 respectively), as well as by the Spanish Agency of Medicines and Medical Devices (AEMPS) – records 858/20/EC and 856/20/EC respectively. The participants volunteered to participate in this study, were informed about the procedures and possible adverse effects, and signed the informed consent to participate.

### 2. Consent for publication

Not applicable

### 3. Availability of data and materials

The datasets used and/or analyzed during the current study are available from the corresponding author on reasonable request.

### 6. Authors’ contributions

LBF, JM, MGS, AMG, FOB and AI conceived and designed the study. LBF, JM, CR and AI developed the electronic hardware and the software. MOK, AS and AI designed and developed the intramuscular electrodes. LBF, JM, CR, MGS, AMG, CRG, FGH, FOB and AI conducted the experiment. LBF and JM performed data analysis and drafted the manuscript. MOK, CR, MGS, AMG, CRG, FGH, AC, AJA, AGA, FG, AS, FOB and AI revised the manuscript critically. All authors read and approved the final manuscript.

## 7. Acknowledgements

The authors would like to express their gratitude to the participants of this study, and to Alejandro Pascual-Valdunciel and Juan Camilo Moreno for their support regarding the documentation for ethical committees and the AEMPS.

